# The diversity of clinical *Mycobacterium abscessus* isolates in morphology, glycopeptidolipids and infection rates in a macrophage model

**DOI:** 10.1101/2024.02.29.582856

**Authors:** Virginia Pichler, Lara Dalkilic, Ghazaleh Shoaib, Tirosh Shapira, Leah Rankine-Wilson, Yves-Marie Boudehen, Joseph Chao, Danielle Sexton, Miguel Prieto, Bradley Quon, Elitza Tocheva, Laurent Kremer, William Hsiao, Yossef Av-Gay

## Abstract

*Mycobacterium abscessus* (Mab) colonies adopt smooth (S) or rough (R) morphotypes, which are linked to the presence or absence of glycopeptidolipids (GPL), respectively. Though clinically relevant, the association between GPL levels, morphotype and pathogenesis are poorly understood. To investigate the degree of correlation between Mab morphology, GPL levels, and infectivity, we generated isolates from Mab-positive sputum samples from cystic fibrosis patients. Isolated strains were categorised based on their morphology, GPL profile, and replication rate in macrophages. Our findings revealed that around 50% of isolates displayed mixed morphologies and GPL analysis confirmed a consistent relationship between GPL content and morphotype was only found in smooth isolates. Across morphotype groups, no differences were observed *in vitro*, yet using a high-content THP-1 cell *ex vivo* infection model, clinical R strains were observed to replicate at higher levels. Moreover, the proportion of infected macrophages was notably higher among clinical R strains compared to their S counterparts at 72 hours post-infection. Clinical variants also infected at significantly higher rates compared to laboratory strains, highlighting the limited translatability of lab strain infection data to clinical contexts. Our study confirmed the general correlation between morphotype and GPL levels in smooth strains yet unveiled more variability within morphotype groups than previously recognised, particularly during intracellular infection. As the rough morphotype is of highest clinical concern, these findings contribute to the expanding knowledge base surrounding Mab infections, offering insights that can steer diagnostic methodologies, and treatment approaches.

## Introduction

Non-tuberculous mycobacteria are a class of saprophytic bacteria represented by over 180 species. Ubiquitously found in the environment (soil, water, vegetation), a subset is classified as opportunistic pathogens for their ability to occupy human niches, primarily as pulmonary or soft tissue infections and frequently affecting immunocompromised individuals [1–4]. The most abundant pathogenic species belong to the Mycobacterium avium complex (MAC; *Mycobacterium avium, Mycobacterium intracellulare*) and Mycobacterium abscessus complex (MABSC: *Mycobacterium abscessus abscessus* (Mab), *Mycobacterium abscessus massiliense* (Mma), *Mycobacterium abscessus bolletii* (Mbo)), the latter which remains an ongoing treatment challenge due to its intrinsic resistance to growing incidence of broader antibiotic resistance [5–6].

Mycobacteria follow a similar pathogenesis strategy, primarily infecting hosts via aerosol inhalation where they interact with and are engulfed by the innate immune cells, namely macrophages, where they survive and replicate in phagocytic vesicles [2, 7–8]. This strategy has been well-documented in Mab infections and is associated with the transition of the bacteria from the smooth (S) to rough (R) morphotype [8]. The difference in intracellular behaviour between the two types has been linked to the presence of glycopeptidolipids (GPL) [2]. Cell surface GPL decorate the S morphotype and mask underlying TLR-2 agonists, such a phosphatidyl-myo-inositol dimannoside (PIM2) [9] and lipoproteins [10] and allow for greater initial colonisation in loner phagosomes. Conversely, R morphotypes are associated with a strong immunological response, persistent infections and presence in social phagosomes.

The colony morphology of the S morphotype has been described as a uniformly round shape, shiny and luxuriant (Figure 1, left), whereas the R morphotype is often irregular in shape, matte, wrinkled, and textured (Figure 1, right). The characteristics of these opposite morphologies has, indeed, been widely described in the literature, however, to our knowledge no deliberate study has been performed to determine a strict set of criteria for each progressive morphotype between the canonical extremes, represented by the S and R laboratory strains ATCC19977/ CIP104536 S/R, denoted as Cip^S^ and Cip^R^ throughout. However, rarely do clinical manifestations perfectly mirror their lab counterparts. This is of importance as morphological transition is a marker of infection progression and establishing these criteria will help reduce inter-observer variability when reporting diagnostic results [11]. Furthermore, following consistent morphological plasticity tracking metrics between clinics and research facilities will help build a more robust pool of data to reveal the full extent of the relationship between morphotype and infection persistence and severity. Herein, we describe the isolation, molecular analysis, and intracellular growth of 46 clinical isolates from cystic fibrosis patients, represented by an approximate equal distribution of the S and R morphotypes.

**Figure 1.**
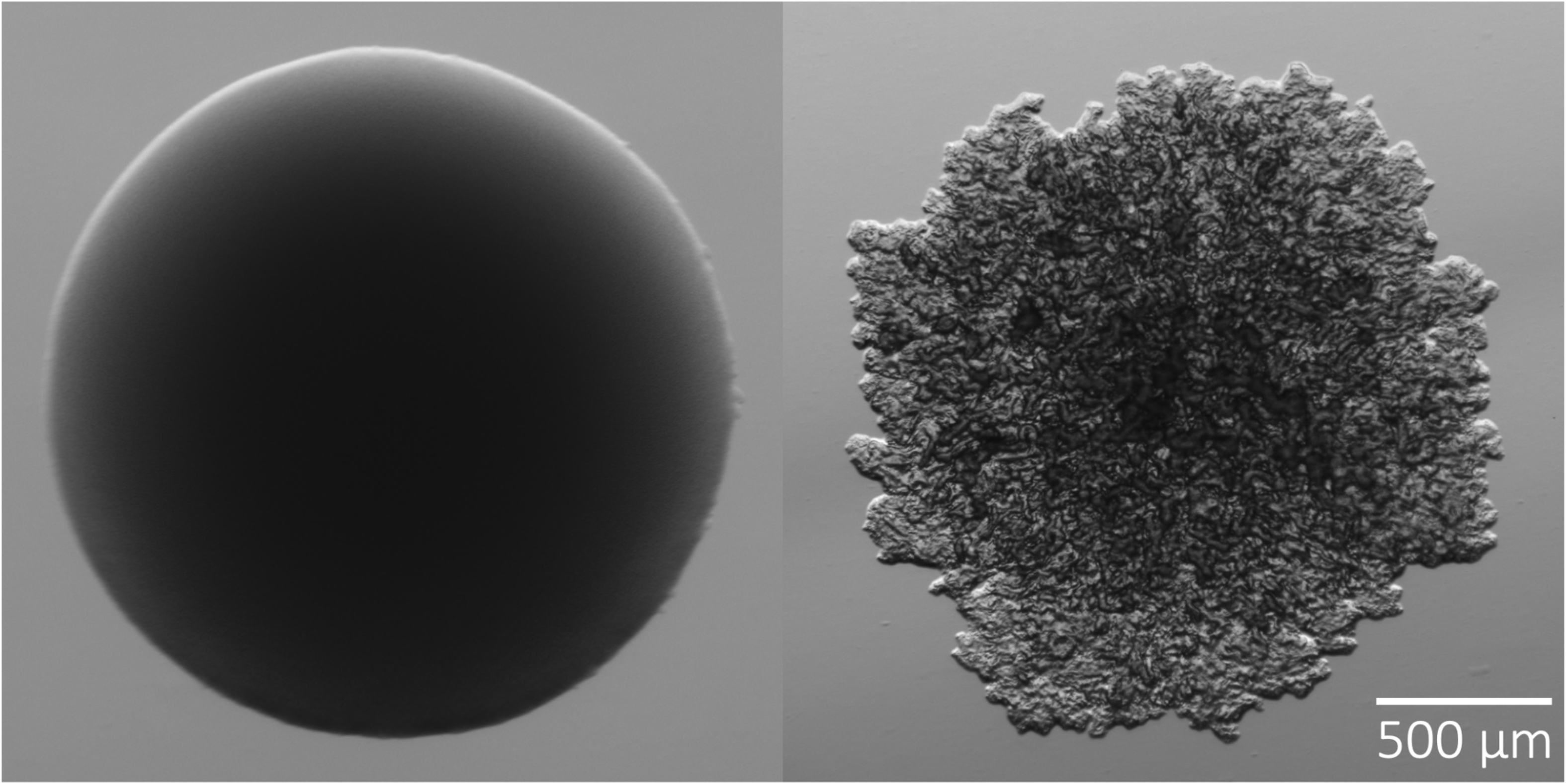
Characteristic two morphotypes of M. abscessus. A. smooth (left) and rough (right). The smooth (S) morphotype is shiny and luxuriant with a uniformly round shape; the rough (R) morphotype is matte, wrinkled, and textured with an irregular in shape.

## Materials and Methods

### Clinical Isolates

This study used a total of 46 laboratory-confirmed mycobacterial isolates that were obtained from 19 CF patients in St Pauls Hospital (Vancouver, Canada). Isolates were confirmed through MALDI-TOF MS and 16s rRNA target sequencing and obtained from the B.C. Centre for Disease Control (BCCDC).

### Strain separation

BCCDC glycerol stocks were streaked onto Luria Bertani (LB) agar plates and incubated at 37 °C for 4-6 days. Isolates were visually inspected for morphotype composition at high resolution with the Zeiss Axio Zoom V16 microscope according to the categorical descriptors in Table 1. Plates containing heterogenous populations were re-streaked until pure, stable morphotypes were achieved. Single colonies were inoculated in 7H9 Broth (Difco™ Middlebrook) supplemented with 10% (v/v) OADC (0.05% oleic acid, 5% bovine albumin fraction, 2% dextrose, 0.004% catalase, and 0.8% sodium chloride solution) and 0.05% (v/v) Tween 80, termed 7H9OADCT throughout. Glycerol stocks of each step were stored at-80 °C.

**Table 1.** Canonical features of smooth (Cip^S^) and rough (Cip^R^) morphotypes.

|  | Cip<sup>S</sup> | Cip<sup>R</sup> |
| --- | --- | --- |
| Appearance | Shiny | Matte, textured |
| Colour | Brown | Gray |
| Margin | Uniform | Irregular |
| Glycopeptidolipid | Present | Absent |
| Infection stage | Colonising | Persistent |
| Antibiotic resistance | Low | High |

### Lipid Extraction

Fresh bacterial lawns on LB agar were harvested and pelleted in glass vials by centrifugation at 2,000*g* for 5 min. Polar lipids (GPL) were extracted using the chloroform-methanol method, as described previously [12]. Final extracts were solubilized in 300 μL of chloroform/methanol (2:1 v/v). Lipids were visualised with thin layer chromatography (TLC) by adding 20 μL of sample on a silica gel 60 F254 sheet (Merck) and placed in a chloroform/methanol/water (90:10:1, v/v/v) solution for migration. An orcinol-sulfuric acid solution (0.2% and 20%, respectively) was sprayed onto the sheet and the plates were lipid profiles revealed using heat. GPL were detected with reference to the control strains (Cip^S^, CIP104536T (S) and Cip^R^, CIP104536T (R)). Replicates were performed for strains where morphotype categorisation did not align with the TLC results.

### Ex-vivo THP-1 macrophage infection model

#### Construction of an integrative vector expressing a fluorescent marker

Derivative versions of the integrative plasmid pMV306 [13–14] were constructed to allow strong expression of either mScarlet (pMV306-mScarlet, Figure S1) or mWasabi (pMV306-mWasabi) fluorescent proteins with a kanamycin resistant cassette for selection. This plasmid contains pMV306 integrase *attP* as a one-step integration construct. Briefly, the mScarlet or mWasabi coding sequences, placed under the control of the constitutive P left* promoter, were PCR-amplified by Q5® High-Fidelity DNA Polymerase (New England Biolabs) using plasmid L5 attB::Pleft*mScarlet/mWasabi (Addgene plasmids 169410 and 169409, respectively) as DNA templates [15]. Primer pairs, dual forward primer P*: ATCTTTAAATCTAGATGGCCGCGGTACCAGATCTT, mScarlet reverse primer: AGCTGGATCCATGGATTCACTTGTACAGCTCGTCCATGCC, mWasabi reverse primer: AGCTGGATCCATGGATTTACTTGTACAGCTCGTCCATGCC. The linear fragments were purified on agarose gels (NucleoSpin Gel and PCR Clean-up, Macherey-Nagel). Following the manufacturer’s instructions, Pro Ligation-Free Cloning Kit (abm®) reactions were performed to insert these linear fragments into the pMV306 previously digested with EcoRV and transformed into *Escherichia coli* (*E. coli*) Stellar TM competent cells (Takara Bio), purified (NucleoSpin Plasmid, Macherey-Nagel) and verified by DNA sequencing. Plasmids were then electroporated into the *M. abscessus* strains [16–17] and recombinant clones harbouring the constructs inserted at the attL5 insertion site in the glyV tRNA gene were selected for on 7H10 agar supplemented with 0.5% glycerol, 10% OADC, and 50 μg/ml kanamycin (7H10OADCKan).

#### Bacterial and THP-1 Cell Culturing

Unless otherwise stated, all bacterial strains were transformed with pMV306-mScarlet and routinely grown from stock at 37 °C in 7H9OADCT with 50 μg/ml kanamycin, termed 7H9OADCTKan. THP-1 human monocyte-derived macrophage-like cells (ATCC TIB-202) cells were grown in incomplete RPMI1640 medium (10% FBS, 2% glutamine and 1% non-essential amino acids) at 37 °C with 5% carbon dioxide (CO_2_). Cell density was kept between 0.25 and 1 × 10^6^ cells/mL for a maximum of three months.

#### Infection

THP-1 cell suspension at 5 x 10^5^ cells/mL in incomplete RPMI1640 was seeded into a 96-well plate at 50,000 cells/well and differentiated into a macrophage-like state over a period of 48 hrs with phorbol12-myristate13-acetate (PMA, 40 ng/mL) at 37 °C with 5% CO_2_. Liquid bacterial cultures were grown to mid-log phase in 100 μL per well in a round-bottomed 96-well plate (VWR), centrifuged (5,000*g*, 10 min) and washed once with sterile Dulbecco’s phosphate-buffered saline (DPBS, Gibco). Cultures were de-clumped using a 25G blunt needle, transferred to a flat-bottomed 96-well plate (VWR) and OD_600_ was measured (OD_600_ of 1 ≈ 1.13 × 10^9^ CFU/mL) using the Varioskan LUX™ microplate reader (Thermo Fisher Scientific). Inoculums of a multiplicity of infection (MOI) of 2 bacteria per macrophage was prepared in incomplete RPMI and opsonised with 10% non-decomplemented human serum for 30 min at 37 °C, shaking. Differentiated macrophages were washed once with warm DPBS and inoculated with the opsonized bacterial suspensions for three hrs (37 °C, 5% CO_2_). Wells were washed twice with DPBS and treated with amikacin (250 μg/mL) for one hr to remove any extracellular bacteria. The amikacin was removed, and wells were washed a final time with DPBS and replaced with 100 μL incomplete RPMI containing 1 μg/mL Hoechst 33342. Plates were incubated for three days at 5% CO2 at 37 °C. Testing of each strain was performed in technical triplicates.

#### High-content intracellular growth analysis

Intracellular bacterial growth was monitored using the CellInsight^TM^ CX5 High Content Platform (Thermo Fisher Scientific) using methods previously described by our group [18–19]. Briefly, macrophages were identified and counted through nuclei staining and a mask was created to represent the entire cell, or region of interest (ROI/circle). Red channel fluorescence (569/593 nm) was used to detect bacteria “spots” inside the cellular ROI. These spots were quantified using a variety of measurements including intensity and area using the Thermo Fisher ScientificTM HCS StudioTM Cell Analysis Software. These fluorescent measurements closely correspond with colony-forming units (CFU), as previously validated [20]. Data were potted using GraphPad Prism version 10 (GraphPad Software, Boston, Massachusetts USA, www.graphpad.com). Measurements were captured at multiple time points to monitor replicate rate and bacterial burden. Super-resolution fluorescence light microscopy of the 62S and 62R isolates expressing mScarlet was used to determine morphotype differences after 72 hrs of infection: macrophages were differentiated onto coverslips and infection was carried out as described above, after which, cells were fixed with 4% paraformaldehyde and imaged using Zeiss LSM 900 confocal microscope equipped with an Airyscan 2 detector and a Colibri 5 light source. Images were collected with a Plan-Apochromat 100x/1.46 oil objective lens and processed using Zen Blue 2.4 software.

## Results

### Subspecies distribution in CF isolates is predominantly Mab

19 patients provided 43 unique sputum samples, which were characterised by the BCCDC into 46 clinical isolates, and further categorized in this study, resulting in 75 individually characterised clinical morphotype strains. Subspecies representation was dominated by Mab in categories such as prevalence within patients (n=14, 73.68%), and total clinical isolates (n=38, 82.61%), (Table 2). Mma was the next most prevalent, with the subspecies isolated in 5 of the 19 patients (26.32%). Mbo was isolated from the single sputum sample provided by patient 16. Smooth and rough morphotypes were evenly distributed amongst patients, isolates and strains. 18 patients returned both smooth and rough morphotypes in their sputum samples, and of the 46 BCCDC isolates collected, 27 (58.7%) were further separated into defined both smooth and rough morphotype strains. All BCCDC isolates were correctly identified to the subspecies level.

**Table 2.**
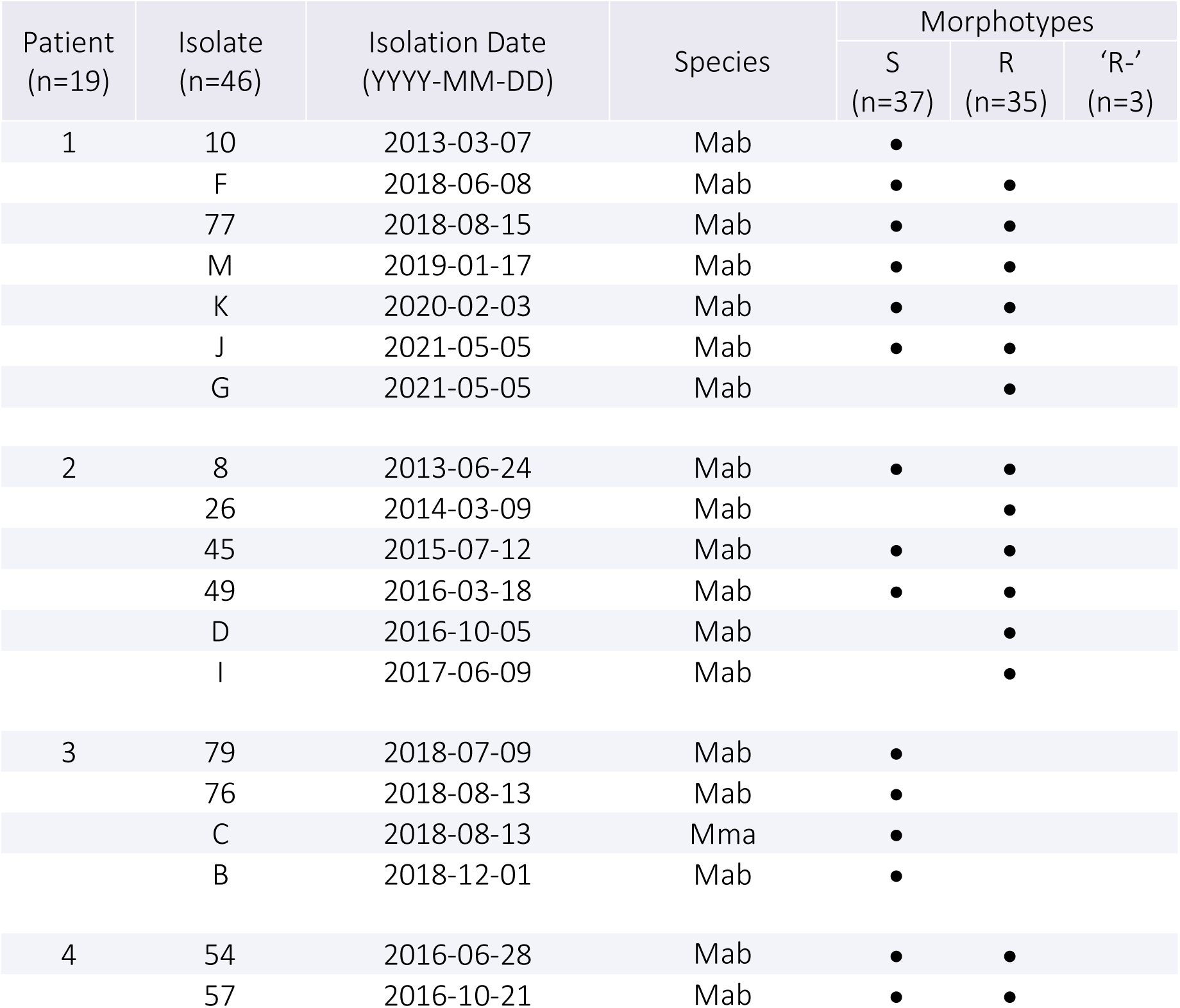
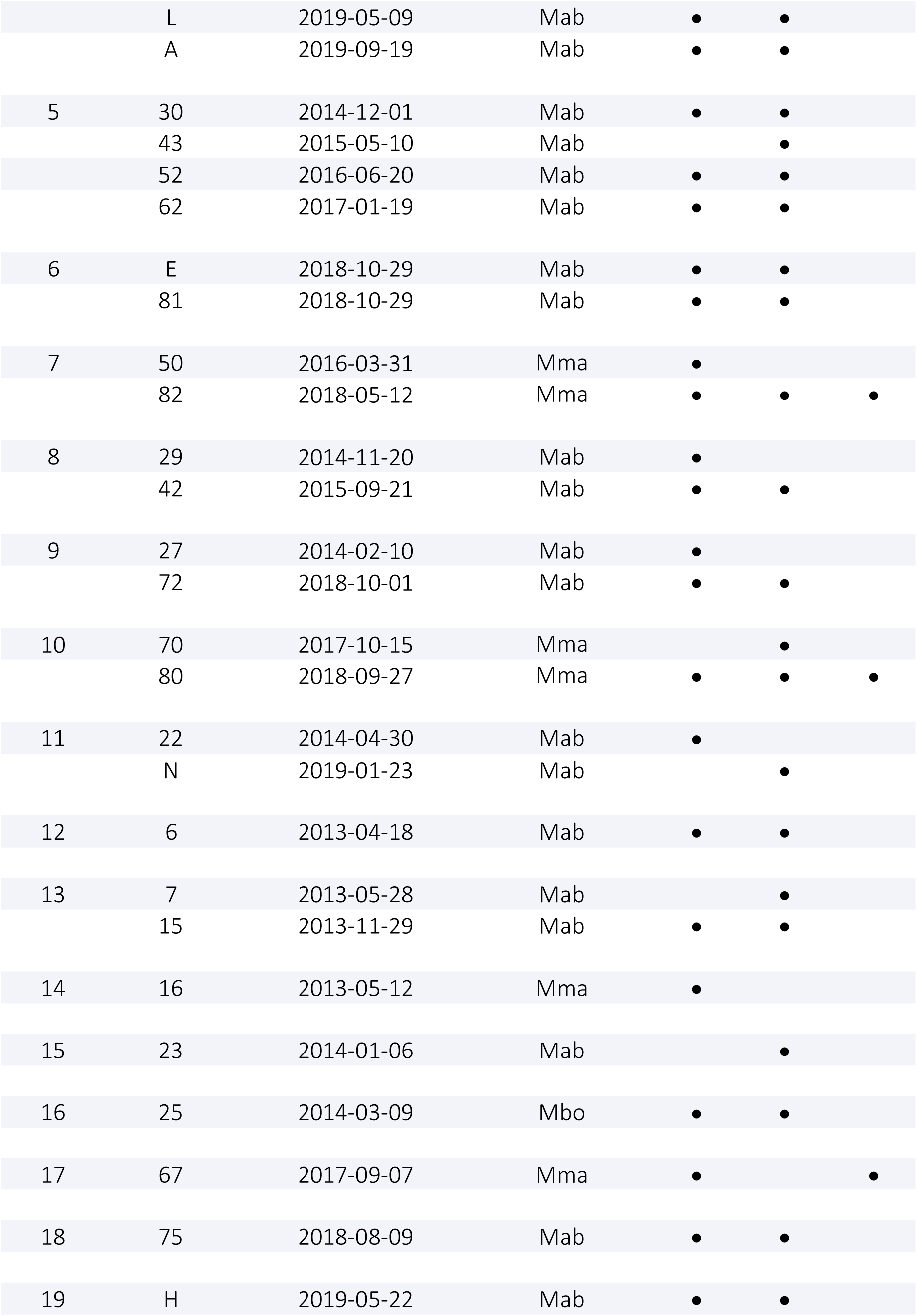
Patient sample isolation dates, species and morphotype categorisation.

| Patient<br>(n=19) | Isolate<br>(n=46) | Isolation Date<br>(YYYY-MM-DD) | Species | Morphotypes |  |  |
| --- | --- | --- | --- | --- | --- | --- |
|  |  |  |  | S<br>(n=37) | R<br>(n=35) | 'R-'<br>(n=3) |
| 1 | 10 | 2013-03-07 | Mab | • |  |  |
|  | F | 2018-06-08 | Mab | • | • |  |
|  | 77 | 2018-08-15 | Mab | • | • |  |
|  | M | 2019-01-17 | Mab | • | • |  |
|  | K | 2020-02-03 | Mab | • | • |  |
|  | J | 2021-05-05 | Mab | • | • |  |
|  | G | 2021-05-05 | Mab |  | • |  |
| 2 | 8 | 2013-06-24 | Mab | • | • |  |
|  | 26 | 2014-03-09 | Mab |  | • |  |
|  | 45 | 2015-07-12 | Mab | • | • |  |
|  | 49 | 2016-03-18 | Mab | • | • |  |
|  | D | 2016-10-05 | Mab |  | • |  |
|  | I | 2017-06-09 | Mab |  | • |  |
| 3 | 79 | 2018-07-09 | Mab | • |  |  |
|  | 76 | 2018-08-13 | Mab | • |  |  |
|  | C | 2018-08-13 | Mma | • |  |  |
|  | B | 2018-12-01 | Mab | • |  |  |
| 4 | 54 | 2016-06-28 | Mab | • | • |  |
|  | 57 | 2016-10-21 | Mab | • | • |  |
|  | L | 2019-05-09 | Mab | • | • |  |
|  | A | 2019-09-19 | Mab | • | • |  |
| 5 | 30 | 2014-12-01 | Mab | • | • |  |
|  | 43 | 2015-05-10 | Mab |  | • |  |
|  | 52 | 2016-06-20 | Mab | • | • |  |
|  | 62 | 2017-01-19 | Mab | • | • |  |
| 6 | E | 2018-10-29 | Mab | • | • |  |
|  | 81 | 2018-10-29 | Mab | • | • |  |
| 7 | 50 | 2016-03-31 | Mma | • |  |  |
|  | 82 | 2018-05-12 | Mma | • | • | • |
| 8 | 29 | 2014-11-20 | Mab | • |  |  |
|  | 42 | 2015-09-21 | Mab | • | • |  |
| 9 | 27 | 2014-02-10 | Mab | • |  |  |
|  | 72 | 2018-10-01 | Mab | • | • |  |
| 10 | 70 | 2017-10-15 | Mma |  | • |  |
|  | 80 | 2018-09-27 | Mma | • | • | • |
| 11 | 22 | 2014-04-30 | Mab | • |  |  |
|  | N | 2019-01-23 | Mab |  | • |  |
| 12 | 6 | 2013-04-18 | Mab | • | • |  |
| 13 | 7 | 2013-05-28 | Mab |  | • |  |
|  | 15 | 2013-11-29 | Mab | • | • |  |
| 14 | 16 | 2013-05-12 | Mma | • |  |  |
| 15 | 23 | 2014-01-06 | Mab |  | • |  |
| 16 | 25 | 2014-03-09 | Mbo | • | • |  |
| 17 | 67 | 2017-09-07 | Mma | • |  | • |
| 18 | 75 | 2018-08-09 | Mab | • | • |  |
| 19 | H | 2019-05-22 | Mab | • | • |  |

### Colony morphology variation among Mab clinical isolates

Colonies were isolated and subcultured several passages to ensure pure morphotypes were reproducible and stable. Strain colonies were inspected using high resolution zoom microscopy and evaluated for morphotype composition. Colonies most often presented as having uniform margins and smooth texture (smooth, “S”, Figure 1, left) or irregular margins with rough texture throughout (rough, “R”, Figure 1, right) and were classified accordingly.

### Visual subcategorization of smooth-like colonies lacks reproducibility

In addition to the canonical morphotypes, several strains appeared as ‘intermediates’, along a spectrum between smooth and rough (Figure 2A). A sub-morphotype classification system was initially used to categorise these intermediates, by way of “S-”, “S--” and “R-“, where S- (Figure 2A, *ii-iv*), is more similar to the reference S morphotype (Figure 2A, *i*), than S--, which displays further edge ruffling and is more matte in appearance (Figure 2A, *v-vi*). This categorization strategy became unwieldy, in that there was no definitive characteristic that would separate any one subcategory from their neighbour, and the variability of shape and texture extended beyond these three discrete groupings. In addition, the intermediate colony morphology strains were often not stably reproducible after repeated subculturing. This phenomenon was observed with both intermediate transitioning to smooth, and vice versa over time. We did not observe, nor subsequently explicitly test for morphotype transition to true rough, and later recategorized S- and S-- into the recognised S morphotype class.

**Figure 2.**
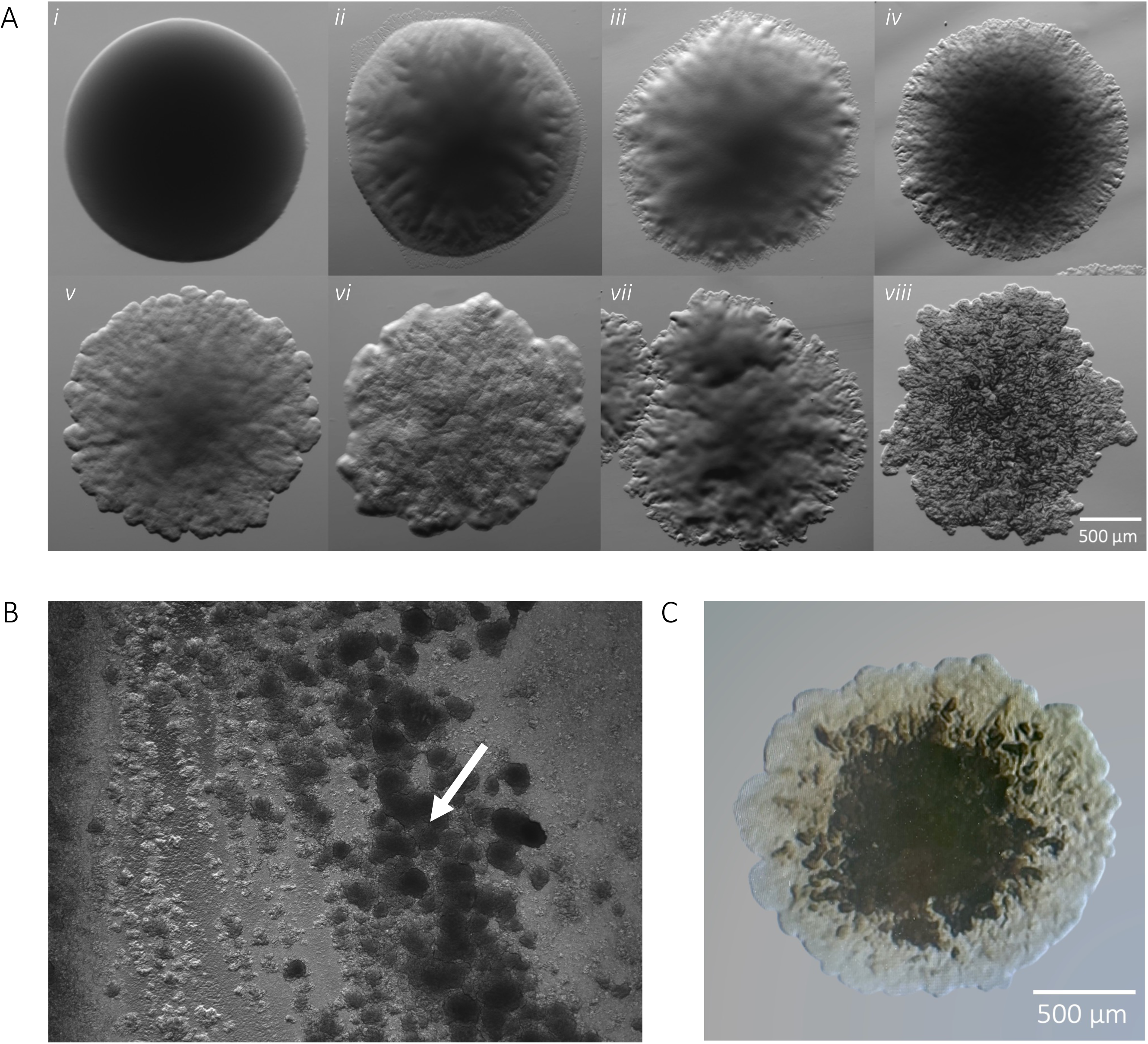
A. M. abscessus morphology variability. Several ‘intermediates’ colony subtypes (*ii-vii*) appear among clinical strains and deviate from the strict smooth or rough morphotype, represented by the first and last images in the series. B. The putative ‘R-‘ phenotype appears rough in individual colonies but adopts a smooth-like characteristic when plated densely (white arrow). C. Putative ‘R-‘ colony with rugged edges, smoother centre and brown colour.

### Some Mab massiliense isolates exhibit density-dependent intermediate morphology

The putative ‘R-‘ category was defined as individual colonies that consistently carried rough-like features, yet when plated densely, would harbour a central, smooth-like phenotype with rough edges (Figures 2B/C). This phenotype was observed in three patients’ unique sputum samples and always belonged to Mma subspecies. 12 (16%) of the 75 strains belonged to Mma. Patient 10 provided two individual samples, one year apart, both only containing Mma (Table 2). The first sample yielded a strict R morphotype Mma (70R), while the second sample contained all three morphotypes (S=80S, R=80R, R-=80R-). To further characterise these morphotypes, we next experimentally assessed lipid composition in each strain to determine the presence of GPL.

### GPL lipid profiles correlate with colony morphology in Mab more than Mma

GPL presence was qualitatively assessed using thin layer chromatography (TLC) of lipid extracts. This assay is a simple way to score strains as GPL-positive or -negative by the presence or absence of banding, respectively (Figure 3A). GPL presence and strain morphotype were correctly matched in 100% Mbo and S typed strains and ∼97% of Mabs isolates (unmatched n=1, J.R). We recorded five patients who provided samples that contained Mma, which were categorised by BCCDC into seven (15.2%) isolates, that we further classified into 12 distinct strains. As previously stated, three of these samples were identified as putative R-morphotype (67R-, 80R- & 82R-), all of which displayed clear banding in the GLP region of the TLC assay, indicating presence of glycopeptolipids. (Figure 3B-C). Retrospectively, all 12 strains would belong to the smooth morphotype group when considering GPL production.

**Figure 3.**
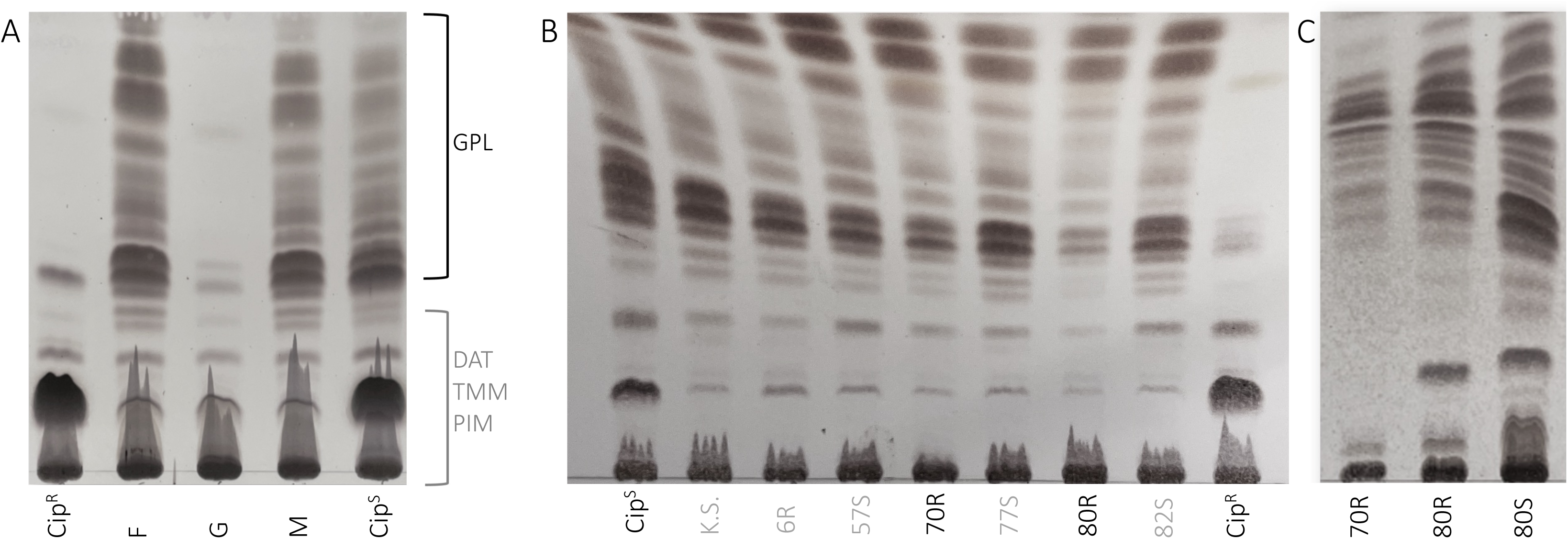
Glycopeptidolipid profiles. A. Cell surface glycopeptidolipids (GPL) are shown in the upper banding of the TLC, with DAT, TMM, and PIM lipids making up the lower bands. GPL present in isolates F and M are smooth morphotypes and match the associated lab strain (Cip^S^); isolate G is a rough morphotype and matches its associated lab strain (Cip^R^). GPL will be present on TLC of mixed morphology isolates, as the rough morphotype is confirmed only by the absence of GPL. Mixed isolates should first be subcultured into pure morphotypes before polar lipid extraction and visualised on a TLC. B. TLC of GPL composition in several clinical isolates, already separated by morphotype. 70R and 80R/R- were morphologically categorised as rough strains yet present with GPL banding. Isolates 70 and 80 are serial collections from the same patient and belong to the subspecies Mma, which may indicate morphological features unique to the subspecies that differ from Mab. C. Repeat extraction and TLC of isolates 70 and 80 to confirm GPL presence.

The strains were subcultured into both solid and liquid media to ensure purity and lipids re-extracted a total of three times, each instance generating consistent results. Furthermore, the GPL profile to morphotype inconsistency was observed in three Mma strains that were grouped as strict R (70R, 80R & 82R). Mma strains 67, 80 and 82 also possessed strict S morphotypes, showing that Mma has more than one morphology associated with the presence of GPL. This leads us to question whether colony morphology should be supported as a method of morphotyping in Mma strains, and further highlights the inconsistency with this method across all MABSC species. We report the GPL production of Mma strains is not readily determined by visual inspection of colony morphology.

### High-content evaluation reveals intracellular replication is highest among clinical R strains

A THP-1 macrophage infection model was used to better approximate the behaviour of the separated strains in the context of host. A total of 69 strains, including reference strains, were successfully electrotransformed with the pMV306-mScarlet integrative plasmid containing the red fluorescent protein, which was used to identify intracellular bacteria burden over time. Relative fluorescence from the transformed bacteria were tracked over three days to assess intracellular replication rate. All fluorescence values were normalised to the first time point at 4 hrs to account for any variations in the MOI. Intracellular bacterial burden was evaluated using several parameters, including the relative replication rate of each strain to its respective lab strain, degree of variance between replicates, and the comparison of each morphotype by collapsing data from all strains into S, R, or R-categories.

### Growth characteristics of the R-subtype is more closely related to S morphotype than R

To evaluate each clinical strain relative to its respective lab strain (Cip^R^ or Cip^S^), a multiple comparisons test was performed with a one-way ANOVA on relative fluorescence captured at multiple time points (Figure 4A) and presented as individual, isogenic pairwise and grouped (Figure 4B, C and D, respectively). Cip^S^ replicated at a rate approximately six-fold greater than Cip^R^ and was found to be statistically significant (p<0.001). Interestingly, clinical strains replicated faster than their cognate lab strain, a characteristic not often reported in clinical strain research. When grouped as isogenic pairs, faster growth was also observed in most R strains compared S strains in the macrophage model (Figures 4C). Indeed, when looking at isogenic strain pairs, nearly all R strains replicated about twice as fast as their S counterparts. Of these, isolates 62, 77, M, A, K and 30 were found to be significantly different. Isolates 77, M and K are sequential collections from Patient 1 and possess both S and R morphotypes. These strains had similar replication rates which were significantly greater (p<0.01) than that of the isolate (FR) collected two months prior, indicating a degree of Mab population stability over the subsequent 18 months of infection (Table 2). Isolates 30 and 62 were the bookends of serial samples provided by Patient 5 for this study and also returned significance between the S and R replication rates, though no significant differences between the other isolates of Patient 5 were found. The replication rate of *in vitro* growth was also measured by OD_600_ and revealed negligible differences between S and R morphotypes, after 72 hours (Figure 4E).

**Figure 4.**
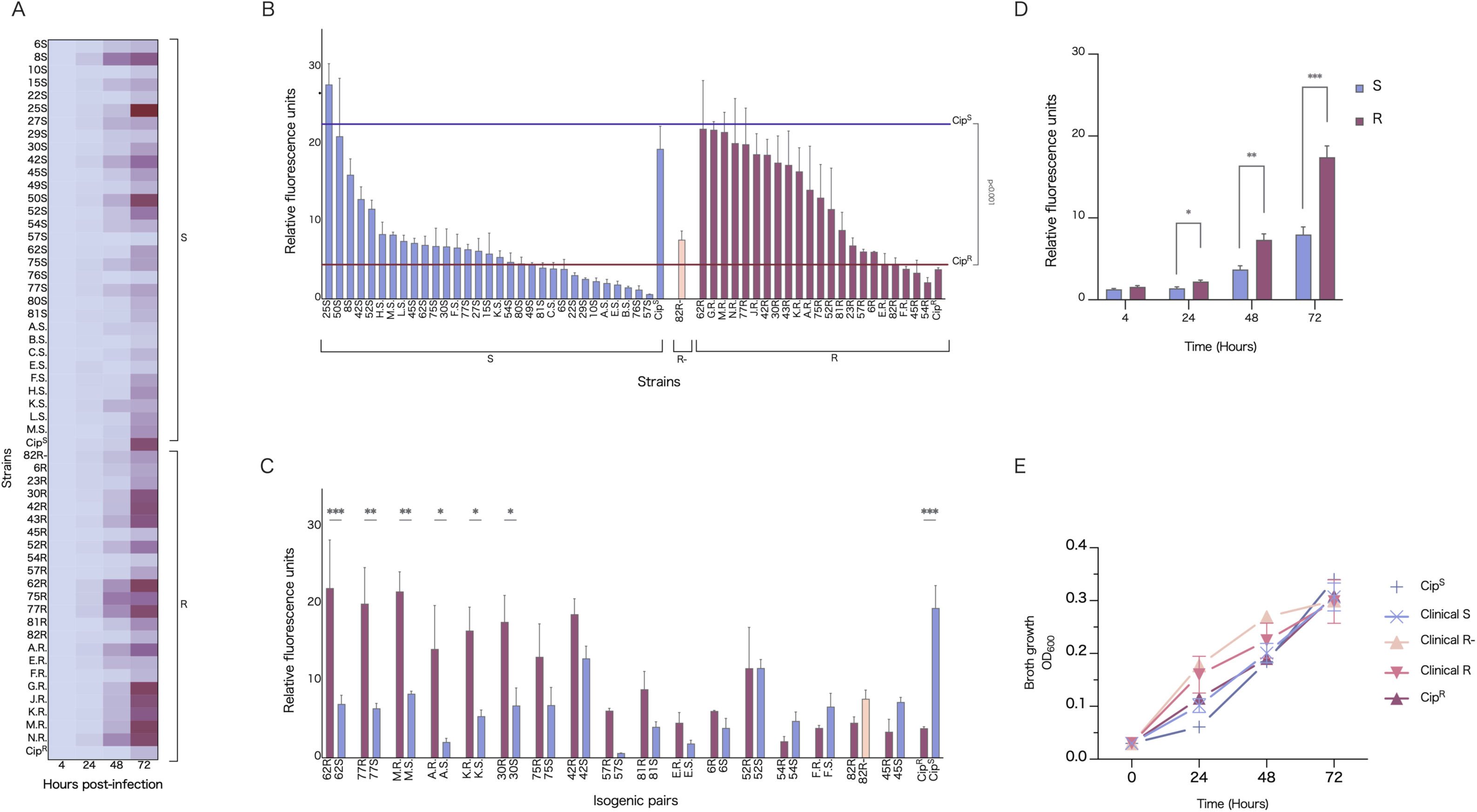
Clinical strain growth rates. A. Relative fluorescence intensity of intracellular M. abscessus clinical strains at 24, 48, and 72 hrs, as normalised to the 4-hr starting time point. All strains were prepared to the same starting inoculum concentration and left to infect THP-1 macrophages for 3 hrs in triplicate, followed by a 1-hr 250 μg/mL amikacin treatment to clear any remaining extracellular bacteria. Lower fluorescence intensity is represented in pale green and high intensity in dark blue. Broadly, S strains (left of plot) replicate as a slower rate than the R strains (right of plot), with the greater divergence appearing at 48 and 72 hrs. Reference strains Cip^S^ and Cip^R^ adopt the inverse replication rate to their clinical counterparts. Overall, there is greater replication rate variability among the R strains. Isolate 82R-, which was classified as an ‘intermediate’ morphotype clusters more closely with the S strains. Other notable outliers, including 25S and 50S belong to the subspecies Mbo and Mma, respectively. B. Intracellular bacterial growth of all isolates at 72 hrs post-infection, grouped by variant. C. Intracellular bacterial growth of isogenic pairs at 72 hrs post-infection. A one-way ANOVA showed statistical significance between the lab strains, Cip^S^ and Cip^R^ (p = 0.0016), and additionally between the isogenic pairs of 62 (p = 0.009), 77 (p = 0.038) and M (p = 0.0052). D. All strains collapsed into morphotype group over 72 hrs. There were no significance differences in rate between groups (S, n=37; R, n=35; R-, n=3). E. Clinical isolate intracellular bacterial replication measured through fluorescence intensity, collapsed by morphotype (S, n=31; R n=22; R-, n=1). R strains replicate at a rate approximately 2-fold higher than that of the S strains and differ significantly at 24, 48 and 72 hrs p.i. The ‘R-‘ intermediate strain (n =1) significantly differed from the R strains, but not the S, indicating it clusters with the intracellular replication behaviour of the S morphotype.

To investigate morphotypes as a group, all fluorescence data were collapsed into the discrete S, R and R-clusters to evaluate for broader trends, not otherwise discernible from individual pairs (Figure 4D). All R strains collectively showed a significant two-fold growth to the S strains at 24, 48 and 72 hrs showing intracellular growth is indeed higher in R than S. The putative R-strain closely matched the replicative behaviour of the smooth strains and replicated at a significantly slower rate compared to the bona-fide R group.

Considering the faster growth of clinical isolates seen in this study, we wanted to see if variance was affected to a greater extent in clinical strains, and if so, which morphotypes contribute to such variation. We performed a Levene’s test to evaluate i) the degree of variance between replicates within a single strain and ii) if any significant differences existed between strains (data not shown). As expected, variance was greatest at 72 hours across all strains, all though no significant differences were observed between strains (p>0.05).

### Reference strains Cip^S^ and Cip^R^ infect macrophages at different rates compared to their clinical counterparts

The proportion of infected THP-1 macrophages was calculated as a percentage of macrophages with intracellular Mab burden over the total macrophage population per condition (Figure 5A). Cip^S^ infected a significantly greater proportion of cells compared to the clinical S strains. The clinical R strains (n=22) infected a greater percentage of macrophages, both compared to the lab strain (Cip^R^) and the clinical S strains (n=31) at 72 hrs post–infection (p<0.05). The high replication rate of Cip^S^ compared to the clinical S strains complements the higher proportion of macrophages infected. Clinical R strains replicated faster and infected a greater proportion of infected macrophages compared to the Cip^R^, whereas inversely, clinical S strains replicated slower and a lesser proportion of infected macrophages compared to Cip^S^. This highlights a divergence in pathogenesis of the clinical strains from their associate lab strains. No significant loss in THP-1 survival was seen over the course of infection, either between morphotypes or strains. Further microscopic analysis of the variants revealed that indeed a feature of the R morphotype at later time points is the formation of extensive extracellular cording, exceeding the size of macrophages and hampering phagocytosis as a clearance strategy (Figure 5B/C/D).

**Figure 5.**
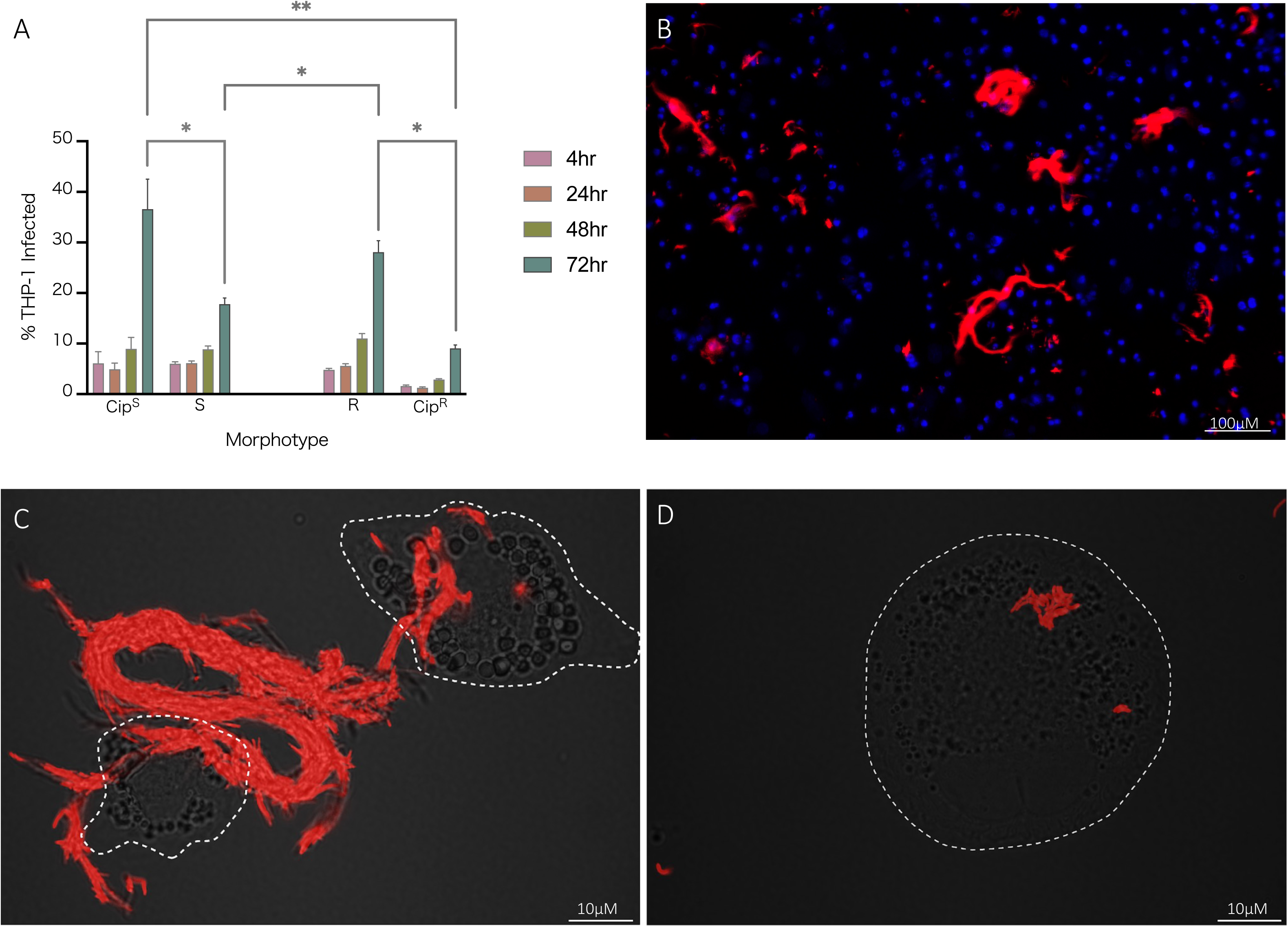
Infection burden. A. Percent of THP-1 macrophage infection by morphotype, clinical and lab strain. The proportion of infected THP-1 macrophages was significantly greater among the R clinical strains compared to their S counterparts at 72 hrs post-infection. Clinical morphotype groups also infected a greater percentage of THP-1 macrophages compared to their associated lab strains. B. High content fluorescent microscopy (CellInsightTM CX5 High Content platform, Thermo Fisher Scientific) image of isolate 62R at 72 hrs post-infection. Macrophage nuclei are stained blue and Mab is expressing mScarlet, with notable cording forming extracellularly. C/D. Super-resolution fluorescence light microscopy of 62R (D) and 62S (E) isolates, 72 hrs post-infection. The R morphotype is found on the surface of macrophages (outlined with white dashed line) with distinct ‘cording’ appearance indicative of cell aggregates too large for clearance by phagocytosis.

## Discussion

This study’s aim to characterise smooth and rough morphotypes of *M. abscessus* clinical infections was initiated with the separation of strains into the two discrete groups. Strict S colonies are characterised by a uniformly round shape, smooth margin, an absence of interior texture and a beige or brown colour, while strict R colonies are characterised by an irregular shape, rough margins, interior texture and a gray colour (Table 1, Figure 1). After multiple passages of each strain to attain sample purity, it became evident that these two morphotype classes were insufficient to capture the full morphological spectrum.

### Colony morphology

Subculturing methods aim to isolate a single, pure, stable sample with only one morphotype present. We found some isolates readily conformed to a stable morphotype, but some were highly variable over more than five passages. It is important to note that subcultures with mixed populations never generated both strict S and R morphotypes on the same plate, but rather contained several sub-morphotypes appearing as ‘intermediates’ between S and R. Initially, these intermediates were classified as ‘S+’, S++’ and ‘R-‘ and fell serially between S and R (i.e., S, S-, S--, R-, R). A subculture isolated from a strict S colony may yield S, S+ and S++ on the same plate, while other strains would only yield more strict S colonies. While the intermediates continued to appear in the subcultures, the inability to successfully isolate consistent sub-morphotypes indicate morphological variations as transient features. Conditions that may have affected colony morphology, such as density, culture age, and media type likely contributed to the observed variability, however, a separate sub-study would need to be performed to properly characterise the full range of factors involved, as has been done on *Pseudomonas aeruginosa* [11]. Despite this limitation, all strains were separated and classified broadly as S and R; all S sub-morphotypes (S+ and S++) fell under the umbrella of S while all strains classified as R adopted the strict morphotype. The ‘R-‘ sub-morphotype, while appearing most like the R morphotype, presented with less texture and brown, rather than gray, in colour (Figure 2C). Additionally, this sub-morphotype forms concentrated, smoothed patches when plated as a lawn (Figure 2B), a feature absent in the strict R morphotype, which is consistently sharply textured even when plated to lawn density. Notably, when reviewing the associated clinical metadata, all strains that possessed the R-sub-morphotype belonged to a congruent subspecies, Mma, and was observed in three of the total seven isolates. While these findings have revealed a common feature of this subspecies’ colony behaviour, the R- sub-morphotype was 1) found in mixed populations, often among strict S and R colonies, and 2) not ubiquitous across all seven Mma strains, and therefore cannot serve as a stable speciation proxy for all Mma isolates with microscopy.

### High-content investigation into MABSC infection dynamics

Clinical *M. abscessus* research has predominantly been performed *in vitro*, which does not adequately reproduce the complexity of the host environment and immunological responses to infection [21]. As the presence of GPL is an important factor in *M. abscessus* pathogenesis, evaluating these bacteria as they interact with macrophages provides a better opportunity to investigate the contributing role of GPL during infection. *For M. abscessus*, a knowledge gap that exists regarding the heterogeneity of a bacterial population and their dynamics during infection may be resolved using high-content analysis. When collapsed by morphotype, we found no significant differences in *in vitro* growth data between groups (Figure 4E). A THP-1 macrophage infection model was then used to better approximate behaviour *ex vivo*. We found inverse relationships between clinical and reference morphotyped strains in their ability to infect a population of THP-1 cells over time (Figure 5A). We also noted clinical R strains grew faster than Cip^R^ in our infection model, a trait that reversed in S morphotypes (Figure 4B). Employing three technical replicates within each experiment was chosen to balance precision and resource efficiency, given that the 54 strains broadly fell into two categories.

Traditional CFU/mL assays include a consistent amikacin treatment to control for extracellular growth, thus ensuring that colony quantification post-lysis exclusively comes from the intracellular environment, however, this ignores the process of bacterial escape from macrophage lysis. Mab infection persistence is a combination of two strategies: 1) intracellular survival and apoptosis suppression by the S morphotype, and 2) bacterial escape and extracellular aggregate colonization (‘cording’) by the R morphotype (Figure 5B/C/D) [1, 8]. By removing the amikacin treatment step over the course of the infection, a truer reflection of infection burden could be seen due to free movement of the bacteria within and between macrophages. Capturing the increase in percent of infected cells demonstrates a degree of bacterial escape and new macrophage infection. Our results show that at 48 and 72 hrs post-infection, the percent of infected macrophages increased in both morphotypes, clinical and lab strains alike. One limitation must be addressed in that our pool of clinical R strains, including most R-strains, was depleted due to difficulty with mScarlet transformation. Some strains underwent several transformation attempts, with additional cell wall perforation methods to enhance DNA uptake [17]. Transformation of mycobacterial species can be especially challenging due to the complex composition of in the cell wall, which often hampers uptake of foreign DNA. The abundance of lipids in the cell wall additionally renders the bacilli hydrophobic, resulting in clumping when grown in media and necessitating the use of de-clumping methods during the preparation of electrocompetent single cells. Rough morphotype strains have been noted as the most challenging to transform and most often form clumps [1]. This was the case for these clinical strains, with ∼88% (n=7) of those unsuccessful belonging to the rough morphotype.

Nevertheless, the greatest increased observed of infected macrophages were observed in the Cip^S^, clinical R, clinical S and finally Cip^R^, reaffirming that the clinical strains do not predictably follow the behaviour of their associated lab strains. Identifying this divergence holds broad implications for our understanding of Mab pathogenesis and persistence. Many studies [1, 7, 22–24] have delved into the importance of morphotype distinction, their unique intracellular behaviours, and their strategies to successfully persist in the host environment, but our findings highlight that the features observed in lab strains may be insufficient to fully appreciate the complete diversity and variability found in clinical strains. Most infection studies to date have been completed using reference strains (ATCC199977/CIP104536S/R) alone or fewer than ten clinical isolates, limiting the translatability of the results to authentic clinical contexts [1, 24–27]. This seems to be especially salient as not only do our results show the breadth of behaviour across the strains during infection but show that clinical morphotypes broadly behave inversely to their lab counterparts.

While the overall trend showed clinical R strains replicating at higher rates than the S strains, large ranges were still observed within each morphotype, with some of the strains more closely matching their associated lab strain. Isogenic strains also varied by fold-difference after 72 hours. 83% of R strains replicated faster than their S counterparts, with 75% of the faster growing R strains replicating at 2-3-fold greater than their S strain pair. Of the pairs that saw S strain outgrow R, we highlight the profiles of Patients 1 and 4, where earlier isolates (F and 54, respectively) showed slower replication in the R versus S. However, this trait was reversed in the isogenic pairs from the next patient collection (77 and 57, respectively). In both instances, this was accompanied by the decline in lung function capacity (data not shown). R colony density and shape variability has been noted by Park et al. (2015) [28] in the context of progressive disease, however, the full phenotype spectrum has yet to be elucidated, as is evident by our findings of varied intracellular replication rate and other data describing mutational and transcriptional changes [27, 29–30]. Strains 25S and 50S displayed the greatest amount of intracellular growth within their morphotype at 72 hrs p.i., however, these two strains belong to the subspecies Mma and Mbo, respectively, which could account for notable difference.

The higher proportion of infected cells seen in clinical R morphotype may be due to bacterial escape and subsequent increased macrophage TLR receptor engagement to immunostimulatory surface molecules, such as PIMs and lipoproteins, that are otherwise masked by GPL in the S morphotype [31]. The stable cell survival between both morphotypes, and increase in infected cells over time do not support the supposition of macrophage apoptosis and bacterial release. The increase in infected macrophages without cell lysis may attributed in part, to the stimulation of tunneling nanotubes (TNTs), membrane channels that connect cells as a rescue function during stress for the transfer of essential cellular components, such as mitochondria [32]. TNT-like structures have been shown to play a role in dissemination of several intracellular pathogens and have been observed in other mycobacteria, including *M. marinum* and *M. bovis* BCG [33–34].

## Conclusions

This study aimed to further characterise the morphological variations and behaviours exhibited within clinical *M. abscessus* infections. We revealed a relationship between clinical and reference strain behaviour in both phenotypic morphotype and intracellular behaviour.

We highlight the complexity of working with clinical isolates, particularly with the heterogeneity of intracellular replication observed even within each morphotype group. By evaluating strain pairs in an intracellular model, we observed R strain enhanced growth rate compared with S, which likely contributed in part to R strain variability. Together, these data encourage the development of specific R and S strain methodologies when assessing host-pathogen relationships.

Considering these findings, we suggested that the existing S-to-R framework should be considered more as a reference point rather than an absolute standard. This framework can serve as a foundation to drive deeper comprehension of infection progression, as well as the development of virulence and resistance mechanisms. Embracing a more flexible approach to morphotype classification could facilitate a more nuanced understanding of *M. abscessus* infections and aid in the development of more effective therapeutic interventions.

## Acknowledgments

The authors wish to thank M. Ko, F. Roquet-Baneres, W. Daher for technical assistance and helpful discussions.

---

**Figure S1.**
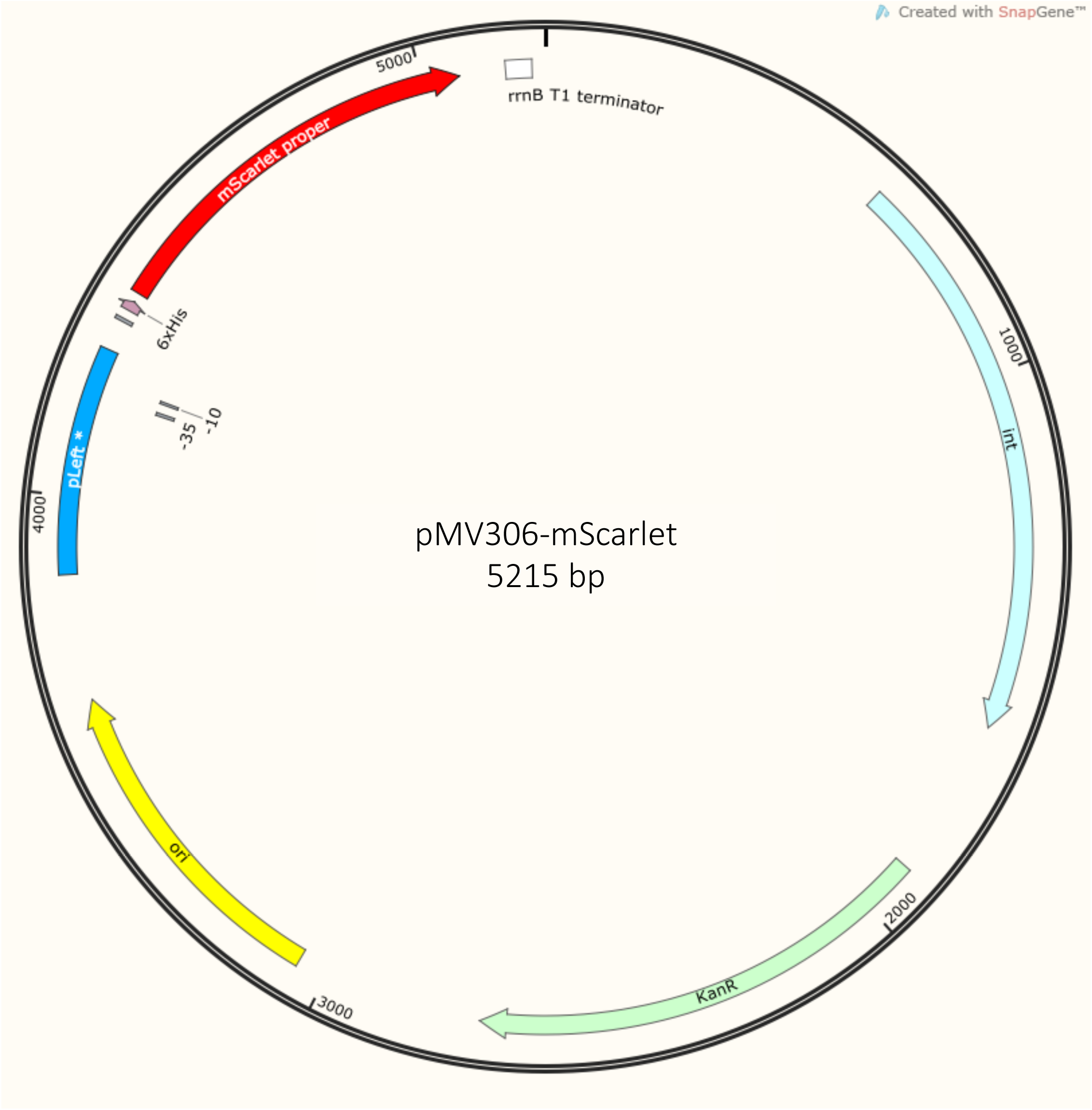
Construct map of the pMV306-mScarlet plasmid.

